# MendelVar: gene prioritization at GWAS loci using phenotypic enrichment of Mendelian disease genes

**DOI:** 10.1101/2020.04.20.050237

**Authors:** Maria K Sobczyk, Tom R Gaunt, Lavinia Paternoster

## Abstract

Gene prioritisation at GWAS loci necessities careful assembly and examination of different types of molecular evidence to arrive at a set of plausible candidates. In many human traits, common small-effect mutations may subtly dysregulate the function of the very same genes which are impacted by rare, large-effect mutations causing Mendelian disease of similar phenotype. However, information on gene-Mendelian disease associations, rare pathogenic mutations driving the disease, and the disease phenotype ontology is dispersed across many data sources and does not integrate easily with enrichment analysis.

MendelVar is a new webserver facilitating transfer of knowledge from Mendelian disease research into interpretation of genetic associations from GWAS of complex traits. MendelVar allows querying of pre-defined or LD-determined genomic intervals against a comprehensive integrated database to find overlap with genes linked to Mendelian disease. Next, MendelVar looks for enrichment of any Human Phenotype Ontology, Disease Ontology and other ontology/pathway terms associated with identified Mendelian genes. In addition, MendelVar provides a list of all overlapping pathogenic and likely pathogenic variants for Mendelian disease sourced from ClinVar.

Inclusion of information obtained from MendelVar in post-GWAS gene annotation pipelines can strengthen the case for causal importance of some genes. Moreover, as genes with Mendelian disease evidence may make for more successful drug targets, this may be particularly useful in drug discovery pipelines. Taking GWAS summary statistics for male-pattern baldness, intelligence and atopic dermatitis, we demonstrate the use of MendelVar in prioritizing candidate genes at these loci which are linked to relevant enriched ontology terms. MendelVar is freely available at https://mendelvar.mrcieu.ac.uk/

## Introduction

The last decade has delivered a bounty of genetic data which has become inexpensive and fast to acquire thanks to advances in high-throughput DNA sequencing and genotyping. On the one hand, this has led to dramatic advances in investigation of the genetic basis of complex, polygenic disease and traits with 8,628 studies featured in the GWAS Catalog as of March 2020 (Buniello et al., 2018). However, the detection of a GWAS signal alone does not identify the causal gene at a locus and so substantial bioinformatics and experimental effort is still required to convert this new genetic knowledge into useful biological insight. At the other end of the spectrum, Mendelian monogenic disease research has benefitted tremendously from recent sequencing methods, which helped to detect and define the causal genes in >1,000 Mendelian conditions (Bamshad, Nickerson, & Chong, 2019). In contrast to complex trait loci, high penetrance, big effect size and typically coding effect of Mendelian perturbations mean the causal gene is more easily detected using statistical methods alone, resulting in a direct link between phenotype and gene. In MendelVar we utilise these direct links between phenotypes and genes from Mendelian traits to aid in identifying causal genes and pathways implicated in GWAS of complex traits.

MendelVar is the first-of-its-kind comprehensive tool linking the information obtained about gene function in Mendelian disease research to help inform candidate gene prioritisation in GWAS. First, it makes it easy to find associations of a given set of genomic intervals or SNPs with Mendelian disease in a range of databases. Secondly, it tests for enrichment of disease phenotype and gene ontology terms associated with detected Mendelian disease. MendelVar is presented as a freely available and regularly updated web-server accessible on https://mendelvar.mrcieu.ac.uk/.

Modern definition of Mendelian disease is complex, with variable levels of monogenicity, penetrance (Claussnitzer et al., 2020), epistasis, gene x environment interactions and indeed pleiotropy observed (Chakravorty & Hegde, 2017). In MendelVar, we follow the working definitions from Chakravorty & Hegde (2017) and Hansen et al. (2019) with broadly defined Mendelian genes to include genes causal for highly penetrant monogenic and oligogenic diseases, including diseases involving cell mosaicism, recurrent *de novo* structural variants causing mostly developmental diseases, such as Smith-Magenis syndrome and some germline cancer susceptibility genes such as BRCA1/BRCA2.

As long as the gene-phenotype association is highly replicable, penetrant, rare, and of large effect, it increases our understanding of biological processes involved in disease pathology and augments our pool for potential therapeutic targets. As such it is included in MendelVar, in line with inclusion in the main reference for knowledge about Mendelian disease – the OMIM database. Still, the majority of relationships presented stick to the strict monogenic Mendelian model (5,524 traits on OMIM, February 2020), followed by susceptibilities to complex disease or infection (694 traits) and somatic cell genetic disease (229 traits).

The number of genes with a known disruption resulting in Mendelian disorder currently totals 4,038 genes in OMIM (February 2020) with ∼300 new Mendelian phenotypes added to the database every year. There is a large scope for an increase in numbers for genes associated with Mendelian disease. Estimates (depending on the type of constraints on the gene) are at between 5,000 to 10,000 potential novel Mendelian disease genes (Bamshad et al., 2019) and proportionately higher number of Mendelian diseases due to pleiotropy (Chong et al., 2015). MendelVar will be regularly updated with new findings on Mendelian disease genes.

It has been long acknowledged that many common complex diseases caused by hundreds of loci with small effect contain a small sub-cohort of individuals with monogenic large-effect disruption in key genes driving complex disease – with examples including coronary artery disease, diabesity, obesity and autism (Chong et al., 2015). Since small-effect mutations in the affected genes circulate in the general at-risk population, function of genes affected in the monogenic forms can inform us about the main biological processes involved in complex disease aetiology and help us develop therapeutics.

It has been hypothesised that many cases involve subtle dysregulation of Mendelian genes’ function, mainly through *cis-* (but also *trans-*) regulatory changes affecting expression, which contribute to risk for phenotypically similar complex disease. Indeed, Freund et al. (2018) have shown that gene sets with confirmed phenotypically-matching or related Mendelian lesions are ∼27 times more likely to be enriched for among all GWAS genes across 62 human traits compared to phenotypically unrelated sets of Mendelian disease genes. Examples include enrichment of growth defect genes in the height GWAS or immune dysregulation genes in a range of inflammatory conditions, such as inflammatory bowel disease. In general, Mendelian disease genes show enrichment among genes flanking the low *p-*value disease GWAS loci and their occurrence positively correlates with association strength (Chong et al., 2015). Widespread comorbidity has also been detected between Mendelian disease and complex disease (Blair et al., 2006) as well as cancer (Melamed, Emmett, Madubata, Rzhetsky, & Rabadan, 2015), which can potentially be driven by pleiotropy (Pividori et al., 2019). The established involvement of certain Mendelian disease genes in complex traits has started to become utilised in evaluating GWAS gene prioritisation algorithms (Barbeira et al., 2019; Guo et al., 2019) and indeed, in gene prioritisation itself (Schlosser et al., 2020). MendelVar aims to simplify this process of integrating information about Mendelian disease to prioritise candidate causal genes at GWAS loci. It has been indicated that Mendelian disease-linked genes make for more successful drug targets (King, Davis, & Degner, 2019; Nelson et al., 2015) and so integration of Mendelian disease data may be especially useful in prioritizing the loci with Mendelian disease evidence for pharmaceutical interventions.

MendelVar allows quick assessment of the likely impact of Mendelian disease-related genes from specified genomic regions (identified by GWAS or other means) on the user’s complex phenotype of interest. It lists the details of all the Mendelian disease genes found in the input genomic intervals, extracted from OMIM and similar databases, as well as closest rare mutations responsible for them available in ClinVar. INRICH is then used for calculating the enrichment of Disease Ontology, Human Phenotype Ontology terms amongst the background of all Mendelian disease-related genes, giving the researcher an overview of any possible shared phenotypic features of identified Mendelian genes with the trait of interest, e.g. in terms of anatomy.

In this paper, we first describe the process of identification, filtering and integration of Mendelian disease data sources. We compare MendelVar with similar tools in terms of analytical features and the breadth of data mined. Following that, we present different possible MendelVar workflows and apply those to dissect top hits from 3 published GWAS studies. MendelVar is shown to allow easy and quick triangulation of Mendelian disease data at the causal gene and enriched biological pathway/phenotype level.

## Results

### Integration of MendelVar data sources

The paramount reference for description of Mendelian disease and their causal genes is the (Online) Mendelian Inheritance In Man (OMIM) database (Amberger, Bocchini, Scott, & Hamosh, 2019). MendelVar uses all the confirmed gene-disease relationships featured in OMIM and complements it with three more specialist data sources for Mendelian disease: Orphanet (INSERM, 1999; Rath et al., 2012) (a database centred on rare, typically monogenic disease), expertly curated gene panels used for diagnostics from Genomics England PanelApp (Martin et al., 2019) and results from on-going Deciphering Developmental Disorders Study (DECIPHER) - whose aim is to identify *de novo* micro genomic rearrangements responsible for undiagnosed developmental delay disorders (Firth et al., 2009).

While OMIM is the established reference for mapping Mendelian disease to genes, adding gene-disease associations from DECIPHER, Orphanet and Genomics England panels allows increased discovery compared to OMIM alone (**Table 1**). The 11,412 gene-disease relationships in MendelVar contain 4,667 unique genes, 629 of them not assigned to any Mendelian disease in OMIM. Integration of the four data sources resulted in 11,412 gene-disease relationships, compared to 6,221 present in OMIM alone. Although there is some redundancy, across Orphanet, DECIPHER and Genomics England, we found at least 1,875 gene-disease associations (either at an individual disease level or same phenotypic series level) that were definitely not in OMIM (i.e. counting only entries that have both gene and disease MIM ID assigned).

**Table 1.** Comparison of MendelVar with its Mendelian disease source databases.

|  | OMIM | Orphanet | Genomics<br>England | DECIPHER | MendelVar |
| --- | --- | --- | --- | --- | --- |
| Number of genes | 4,038 | 4,096 | 3,022 | 1,750 | 4,667 |
| Number of diseases | 5,345 | 3,812 | 4,122 | 2,080 | 6,936 |
| Number of gene-disease associations | 6,221 | 7,605 | 4,679 | 2,292 | 11,412 |

MendelVar includes short disease descriptions sourced from OMIM, Orphanet, Uniprot (Breuza et al., 2016) and Disease Ontology (DO) (Schriml et al., 2019). Out of 11,412 gene-disease entries in the MendelVar database, 9,495 contain a disease description, amounting to 5,878 unique disease descriptions.

In addition, MendelVar cross-references input genomic intervals against ClinVar (Landrum et al., 2019) for pathogenic or likely pathogenic variants implicated in Mendelian disease allowing identification of mostly genic variants and repeats in LD with region of interest. Due to small differences in genome builds, 109,164 and 107,407 ClinVar variants are present in the GRCh37 and GRCh38, respectively. 55,474 and 36,668 unique variants in GRCh38 have been directly assigned to 5,218 and 2,237 OMIM and Orphanet disease identifiers. However, in the majority of cases variant submitters did not include this identifier information and used non-standardised disease names which obscures their link to Mendelian disease entries in OMIM and Orphanet.

Following extraction of Mendelian disease-linked target genes from the MendelVar database, MendelVar provides enrichment testing of any specific phenotypic class associated with the disease in question. This is done by first annotating the target genes with all the ontology terms associated with all the Mendelian diseases caused by any mutation in the gene. MendelVar provides easy enrichment testing using human Disease Ontology (DO) and Human Phenotype Ontology (HPO), their respective slim ontologies and Freund et al. (2018)’s 20 Mendelian gene sets based around broad phenotypic categories. Disease Ontology focuses on defining a hierarchy of disease aetiology classes which include affected anatomical entity, part of metabolism, amongst others. Human Phenotype Ontology’s (Köhler et al., 2019) main remit is classification of clinical symptoms and abnormalities associated with disease.

In MendelVar, HPO terms were collected first from OMIM, and then complemented with Orphanet, Decipher DDG2P and official HPO annotations, which altogether contributed HPO terms to 410 diseases with no previous HPO terms in OMIM. Similarly, DO terms were first mined from OMIM, followed by Orphanet and the official DO annotation. The latter two sources provided DO annotation to 249 diseases with no previous annotation in OMIM. Out of 11,412 gene-disease entries in the MendelVar database, 9,810 (5,542 unique diseases) have at least one HPO term assigned and 7,112 (3,755 unique diseases) have a DO term assigned. In total, we include 8,722 distinct HPO and 3,849 DO terms in the MendelVar database. The mean number of all HPO terms and DO terms is 136.6 (18.6 independent leaf terms) and 9.7 (1.9 independent leaf terms) per disease-annotated gene, accordingly.

MendelVar also incorporates the Gene Ontology (Carbon et al., 2019), Pathway Commons (Cerami et al., 2011), Reactome (Fabregat et al., 2018) and ConsensusPathDb (Herwig, Hardt, Lienhard, & Kamburov, 2016) allowing custom testing for enrichment of molecular and biochemical gene pathways against the null background of all Mendelian disease genes in the human genome.

### User input for MendelVar webserver

The MendelVar webserver accepts a maximum input of 10,000 genomic positions or intervals, with a maximum interval size of 20 Mbp.

MendelVar offers three general routes through the pipeline depending on the type of user input. The user can either submit pre-defined GRCh37 or GRCh38 genomic intervals based on their own analysis pipeline (**Figure 1** *right*) or a list of single genomic positions (e.g. GWAS lead SNPs) from which MendelVar will create the intervals to be used. These intervals can be created in two ways. Either, user-specified left and right flanks can be generated around the input positions, up to a maximum of 10 Mbp in each direction (**Figure 1** *left*), or, flexible LD-based intervals can be created using LDlink LDproxy app (Machiela & Chanock, 2015), either through dbSNP rsIDs or single positional coordinates (**Figure 1** *centre*) from GRCh37 (hg19) or GRCh38 (hg38) assemblies. Using an LD statistic of choice: *r*^*2*^ or *D’*, linkage disequilibrium (LD) is calculated within the 1 Mbp window centred on input SNP with LDlink. Boundaries around the input variant are generated by finding the most distant upstream and downstream variant with a minimum specified LD value (between 0-1) in one of the six target 1000 Genomes populations: CEU (Utah residents of northern European descent), EUR (European), EAS (East Asian), SAS (South Asian), AFR (African), AMR (Ad-mixed American). This generated interval can be extended to the nearest recombination hotspot with recombination rate >3 cM/Mb (Myers, Freeman, Auton, Donnelly, & McVean, 2008) based on HapMap II. As LDlink accepts only GRCh37 coordinates, we initially map user-submitted GRCh38 coordinates to GRCh37 coordinates with the UCSC liftOver tool (Haeussler et al., 2019). Therefore, the last step in the case of user input in GRCh38 coordinates is to convert the LDlink-created GRCh37 intervals, optionally extended to the nearest recombination hotspot, back to GRCh38 coordinates using the UCSC liftOver tool.

**Figure 1.**
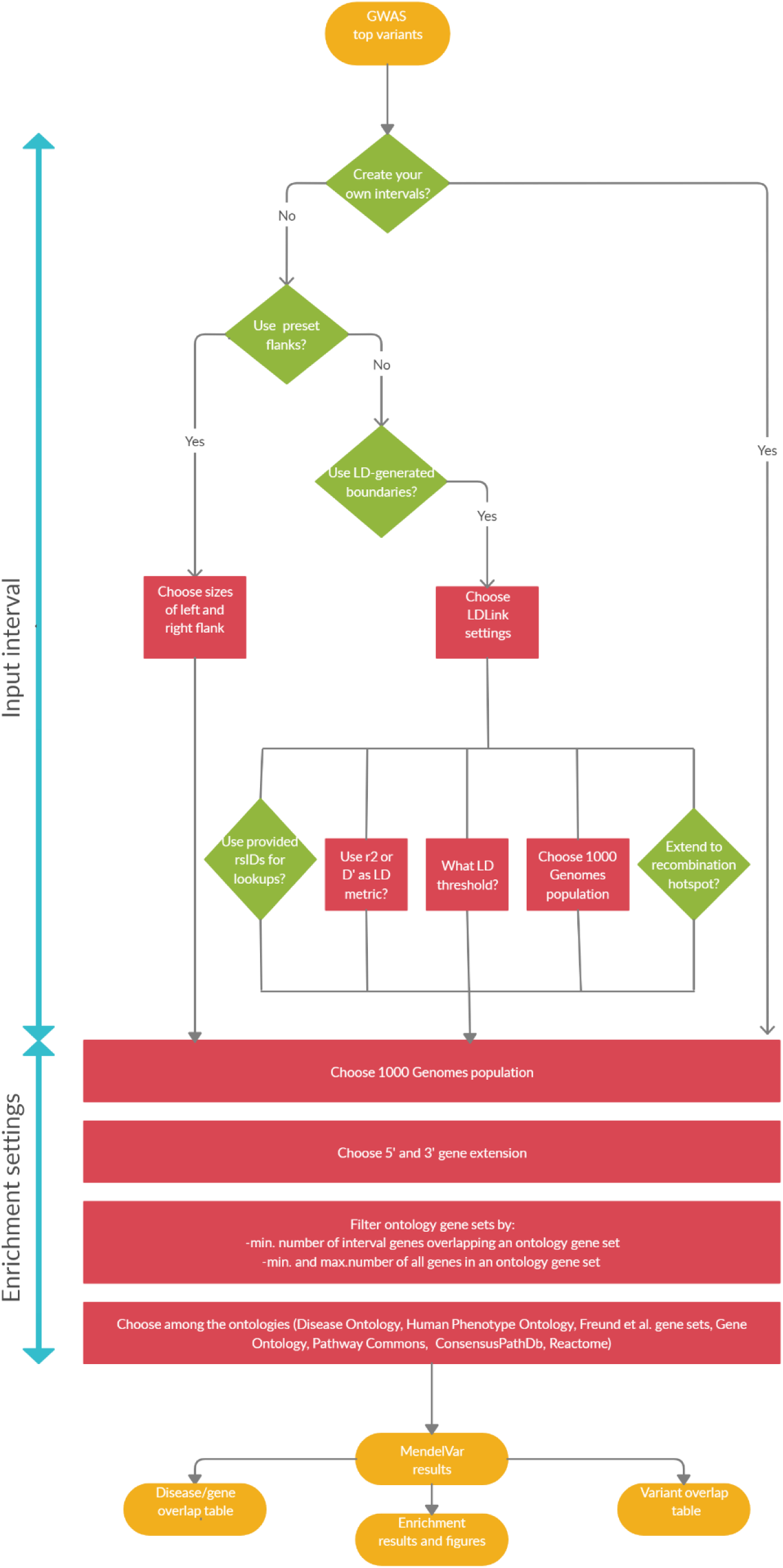
A flowchart demonstrating three possible user routes through MendelVar: a) *left* –MendelVar generates fixed genomic intervals using pre-set left and right flanks against a user-submitted list of genomic positions b) *centre* – MendelVar generates flexible genomic intervals using LD pattern in the region around each user-submitted position/variant rsID. c) *right* – MendelVar accepts user-submitted genomic intervals The genomic intervals generated or obtained from user are subsequently bisected with coordinates for genes and variants known to cause Mendelian disease. Ontology terms associated with Mendelian disease in Human Phenotype Ontology, Disease Ontology are propagated to causal genes and are tested for enrichment among target genes in input genomic intervals. MendelVar also provides an option for enrichment testing with Gene Ontology and biological pathway databases.

### Identification of overlapping Mendelian genes

Genes are defined as overlapping with the genomic intervals based on coordinates of canonical APPRIS (Rodriguez et al., 2013) transcript isoforms if available or alternatively the longest transcript for a given gene; the overlap step is performed with GIGGLE (Layer et al., 2018). Users can extend the gene region by up to 20 kbp in upstream or downstream direction since most of the strong eQTLs (expression quantitative trait loci) regulating gene expression are found in that region (Veyrieras et al., 2008). It is not recommended to extend the gene region too much, as this can result in more genes overlapping each other and being collapsed by INRICH (P. H. Lee, O’Dushlaine, Thomas, & Purcell, 2012), which will then result in a loss of power in the “gene” INRICH mode (*see below*).

### Enrichment testing

Following identification of Mendelian disease-associated genes overlapping the input genomic intervals, MendelVar allows testing for enrichment of terms associated with those genes relative to the background of all Mendelian disease-associated genes in the whole genome. Especially of interest in GWAS for complex traits and uniquely compared to other tools, we allow simultaneous testing using Human Phenotype Ontology, Disease Ontology and Freund *et al*. (2018) gene sets. To give the user an overview of general categories associated with both ontologies and boost statistical power, MendelVar includes the official DO slim (24 terms) and custom-generated HPO slim (25 direct descendants of the root term “Phenotypic abnormality”). In addition, general gene enrichment testing is available with Gene Ontology and its slim, and pathway enrichment testing with ConsensusPathDB, PathwayCommons and Reactome.

Enrichment testing in INRICH can be run in two modes: “gene” and “interval”. In the “gene” mode, the enrichment statistic is calculated over the number of genes in all intervals overlapping a given ontology term. In the “interval” mode, the enrichment statistic is calculated over the number of intervals with at least one gene (regardless of gene number within interval) overlapping a given ontology term. In general, “gene” mode should result in higher power to detect enrichment in the context of typical MendelVar usage, if we expect multiple genes with related functions to cluster in a single locus. The flip side of that in the context of GWAS is that this enrichment value will be biased by the same GWAS signal counted multiple times.

### MendelVar output

Average run time using 100 single variant positions, interval generation with LDlink and enrichment testing using all ontologies is 5 hours due to re-sampling and bootstrapping steps in INRICH. Two main results tables are produced, which detail the overlap of input genomic intervals with Mendelian-disease causing genes and variants, respectively (*see example* **Supplementary Table 1 & 2**). Results of enrichment testing using each ontology are provided in individual tables and are summarised in a set of figures (*see example* **Figure 2 & 3**).

**Figure 2.**
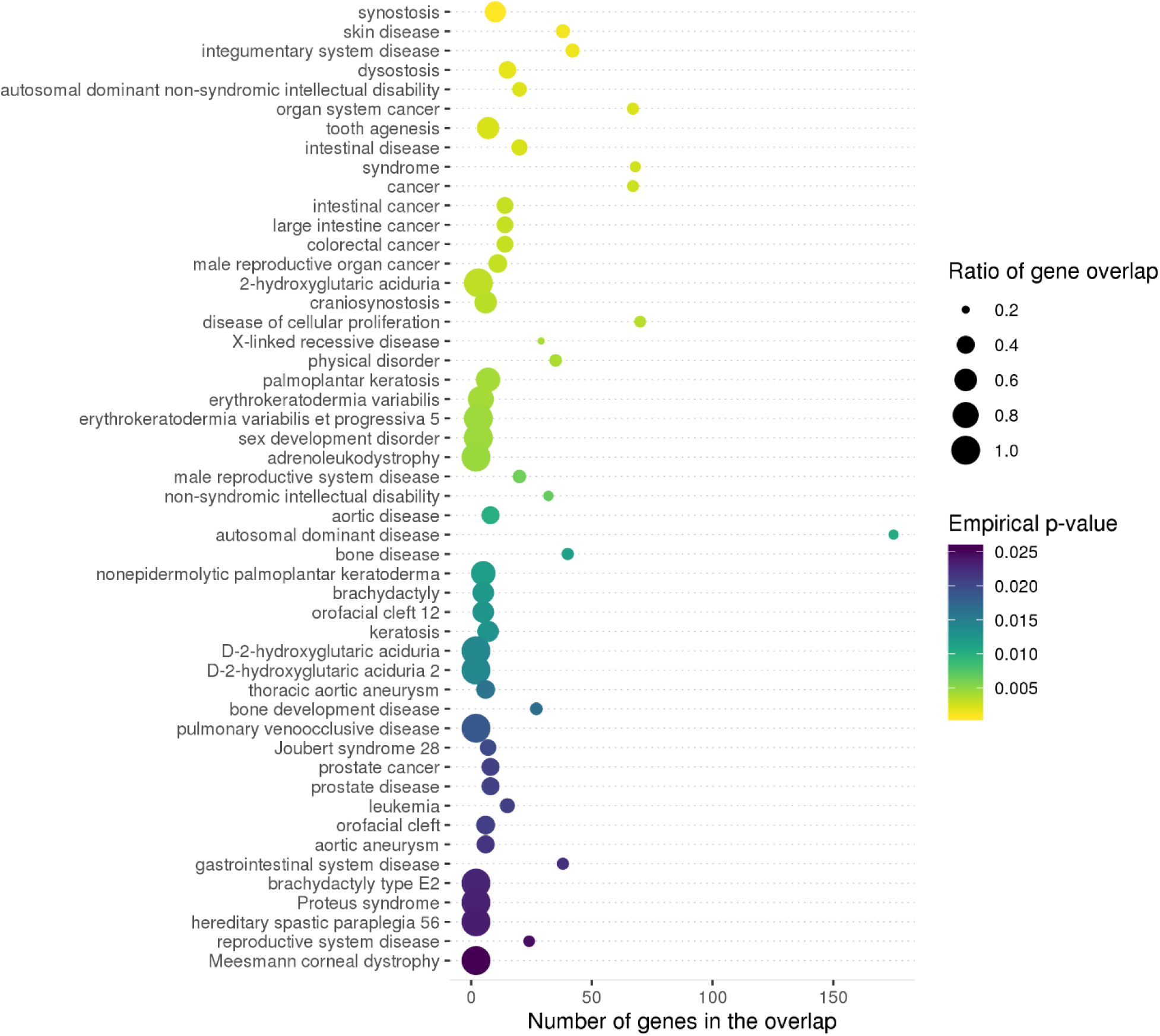
Top enrichment results for Disease Ontology (DO) terms among Mendelian disease genes located within 1 Mbp of lead SNPs in Yap et al. (2018) male pattern baldness (MPB) GWAS. An example of a summary figure produced by MendelVar pipeline depicts top 50 most significant terms in the ontology (*y* axis), sorted first by empirical *p*-value followed by the number of genes overlapping the ontology term in the tested genomic intervals in case of ties (also shown independently on the x axis). The size of each data point represents ratio of gene overlap, i.e. number of genes found in the tested genomic intervals divided by all Mendelian genes annotated with that ontology term available in the genome. Each data point is coloured according to empirical *p*-value for enrichment significance.

**Figure 3.**
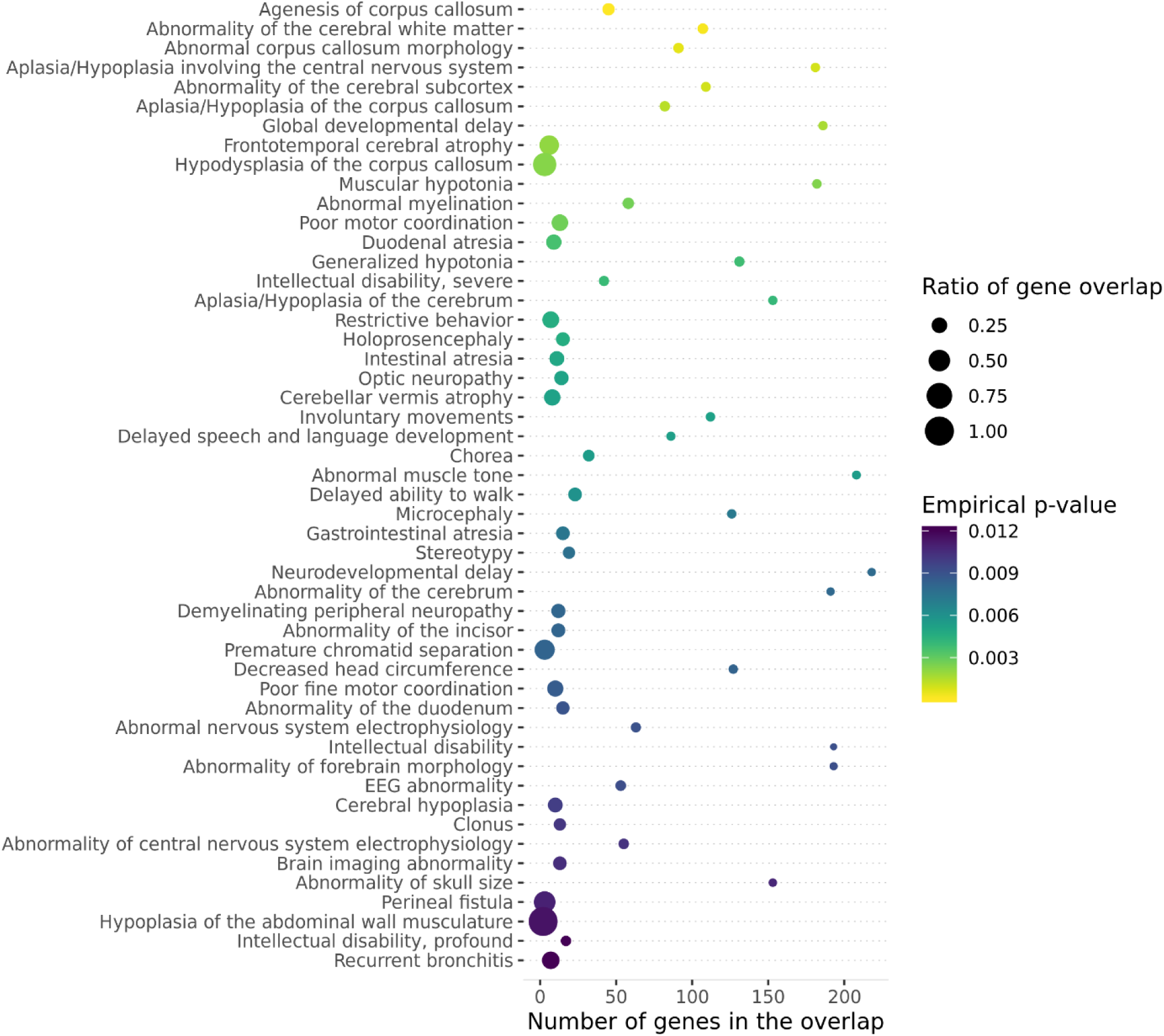
Top enrichment results for Human Phenotype Ontology (HPO) terms among Mendelian disease genes located within 1 Mbp of lead SNPs in Savage et al. (2018) intelligence GWAS.

### Comparison with related tools

We compared MendelVar to 7 most closely matched applications, which also facilitate the use of Mendelian disease data in human genetics research (**Supplementary Table 3**). These tools show differences in accepted inputs, datasets included and overall philosophy and goals of the analysis. Broadly scoped tools such as FUMA (Watanabe, Taskesen, Van Bochoven, & Posthuma, 2017) provide an excellent starting point for GWAS annotation by providing access to rich compendium of resources and analyses, such as gene expression, chromatin interactions, eQTLs. However, by the virtue of being so wide-ranging, they provide only a limited overview of the GWAS link to Mendelian disease and their phenotypes. Packages such as VarfromPDB (Cao et al., 2017), while integrating a selection of the datasets present in MendelVar are built on a reverse premise and mine for variants and genes associated with a phenotype/disease input by the user. Three tools: MARRVEL (Wang et al., 2017), FUMA (Watanabe et al., 2017) and DisGeNET (Piñero et al., 2017), annotate input genes with matching Mendelian disorders from only a subset of databases integrated in MendelVar. None of these 3 tools includes annotation with the full set of human Disease Ontology and Phenotype Ontology terms or features enrichment testing of these terms.

Another comparison group consists of tools such as clusterProfiler (Yu, Wang, Han, & He, 2012), DAVID (Huang, Sherman, & Lempicki, 2009) and Enrichr (Kuleshov et al., 2016). These are popular gene set enrichment packages widely employed in interpreting long gene lists resulting from high-throughput genomics experiments. Despite a large number of compatible ontologies, they never feature more than one of the key ontologies (DO, HPO) available in MendelVar and do not carry out the background gene selection sampling aware of genomic confounders such as SNP density and gene size/density which are required for appropriate GWAS analysis. Simulations have shown that ignoring these confounders in enrichment analysis in GWAS studies and assuming independence can result in up to 100% type I error in cases where it never exceeds the nominal 5% in INRICH (P. H. Lee et al., 2012). Finally, in terms of flexibility of user-input, other than Enrichr, MendelVar is the only enrichment tool to accept genomic coordinates.

We expect that MendelVar users may also utilise other tools in order to triangulate evidence across different data sources, but MendelVar provides the most comprehensive analysis of the relevant Mendelian disease gene data and aims to do so in a flexible and user-friendly way.

### MendelVar use cases

#### I. Yap *et al* (2018) male pattern baldness GWAS

Male pattern baldness (MPB) is a heritable (*h*^*2*^_SNP_ = 0.39) and common condition investigated with GWAS by Yap et al. (2018). They identified 624 near-independent genomic loci, out of which 507 reside in a 1 Mbp window overlapping a Mendelian disease gene.

Results from our gene-based analyses show clear consistency with post-GWAS annotation produced in the original paper using alternative methods (which did not include Mendelian disease phenotype mapping), providing further independent evidence for the prioritization of these genes. Disease Ontology (**Figure 2**) and Human Phenotype Ontology (**Supplementary Figure 1**) analyses in MendelVar support Yap et al (2018)’s conclusion of enrichment of genes involved in mesenchymal lineage specification and odontogenesis as well as positive correlation with bone mineral density (e.g. *integumentary system disease, synostosis, tooth agenesis, hypoplasia of the maxilla* enriched in MendelVar). Similarly, not unexpectedly given the location of hair follicles in the skin, we confirm the importance of genes involved in skin/epidermal development, with the following terms found enriched by MendelVar: *skin disease* and specific subtypes - *(erythro)keratoderma and keratosis, abnormality of skin/nail/eyebrow morphology/hair quantity, sparse eyelashes*. Also, we confirmed pleiotropy of male pattern baldness with reproductive traits and androgen signalling with enrichment of *male reproductive organ cancer, sex development disorder, male reproductive system disease* in the MendelVar results. Pathway enrichment results (**Supplementary Figure 2 & 3**) further highlight *coregulation* of *androgen receptor activity, estrogen-dependent gene expression* and *TGB-β receptor/signalling pathway* which is known to interact with androgen signalling (Pirastu et al., 2017).

An example on how MendelVar can be used to narrow down the list of potential candidate genes is provided at locus 12q13.13 which harbours two lead SNPs (rs3842942, rs7961185) in the MPB GWAS. The SNPs are located upstream of a cluster of 20 keratin (*KRT*) genes. Out of the 20 genes, 17 are annotated with a Mendelian phenotype affecting the integumentary system, however only 6 demonstrate phenotypes affecting hair and scalp (**Table 2**). Three out of the six prioritised genes – *KRT71, KRT74* and *KRT73* are also the top prioritised genes presented in the original paper based on FUMA analysis (Watanabe et al., 2017) and they include the closest gene to index SNPs (*KRT71*).

**Table 2.** Prioritisation of 6 out of 20 proximal keratin genes potentially responsible for two MPB GWAS signals at the 12q13.13 locus using MendelVar annotations from DO and HPO.

| Gene | Relevant ontology terms | Mendelian condition | Prioritise? |
| --- | --- | --- | --- |
| KRT86 | A, B, C, D, F, G | Monilethrix | ✓ |
| KRT81 | A, B, C, D, F, G | Monilethrix | ✓ |
| KRT83 | A, B, C, D, F, G | Monilethrix | ✓ |
| KRT85 | A, B, C | Ectodermal dysplasia 4 | ✓ |
| KRT71 | B, C, D, E, G | Hypotrichosis, wooly hair | ✓ |
| KRT74 | B, C, D, E, G | Hypotrichosis, wooly hair, ectodermal dysplasia 7 | ✓ |

**A** - Patchy alopecia/ Alopecia of the scalp
**B** - Abnormal hair quantity
**C** - Abnormality of hair texture/Fine hair
**D** - Hypotrichosis
**E** - Hair disease (DO)
**F** - Monilethrix (DO)
**G** - Slow-growing hair

#### II. Savage *et al* (2018) intelligence GWAS

Intelligence is a highly complex trait with moderate heritability (*h*^*2*^_SNP_ = 0.19-0.22) which was recently investigated with a GWAS (Savage et al., 2018) that detected globally significant signal at 205 genomic loci. MendelVar analysis highlighted 454 genes causal for Mendelian disease to be positioned within a 1 Mbp window around the loci lead SNPs. 200 genes (43%) identified by MendelVar overlapped 859 candidate genes prioritised by Savage et al. (2018). Comparison of MendelVar enrichment results using Disease Ontology and Human Phenotype Ontology with the alternative post-GWAS gene prioritisation presented in the original paper shows good concordance (**Figure 2, Supplementary Figure 4**). Both analyses reveal enrichment of genes broadly involved in central nervous system development and function; and highlight association with autism and intellectual disability/delay genes (**Table 3**). Out of 114 genes annotated with a “nervous system disease” DO term in MendelVar, 48 (42%) are prioritised among 859 genes highlighted in the original paper. Similarly, 149 out of 326 (45.7%) genes annotated with “abnormality of the nervous system” match those in the Savage et al. results.

**Table 3.** Correspondence of MendelVar DO/HPO enriched terms to the categories highlighted in the Savage et al. (2018) intelligence GWAS paper. Each row represents comparison of MendelVar DO/HPO enriched terms to the related categories highlighted among original gene mapping results in the Savage et al. (2018) intelligence GWAS paper.

| MendelVar enrichment | Savage *et al.* (2018) post-GWAS analysis |
| --- | --- |
| central nervous system disease, congenital nervous system abnormality, abnormal myelination, abnormality of CNS electrophysiology, microcephaly | Mapped genes enriched for:<br>-regulation of nervous system development,<br>-central nervous system neuron differentiation,<br>-regulation of synapse structure or activity |
| frontotemporal cerebral atrophy, abnormality of the cerebral subcortex, abnormality of cerebral white matter, basal ganglia disease, abnormal corpus callosum morphology | Gene *p*-values associated with expression in frontal cortex, striatal medium spiny neurons, hippocampal CA1 pyramidal neurons and cortical somatosensory pyramidal neurons |
| autism spectrum disorder | Positive genetic correlation with autism |
| syndromic intellectual disability, neurodevelopmental delay, delayed speech and language development, delayed ability to walk | Enrichment of mapped genes linked to intellectual disability and developmental delay |

#### III. Paternoster *et al*. (2015) atopic dermatitis GWAS

Lastly, we believe that MendelVar could be potentially useful for mining genes whose Mendelian phenotype matches the one exhibited in our complex disease of interest. This could be especially useful in investigating sub-significant GWAS loci, to determine which loci (of all those that reach a suggestive *p*-value threshold) also have consistent independent evidence implicating loci gene with the disease. As an example, we investigated 119 loci with *p*-value of between 5 × 10^−8^ to 1 × 10^−4^ in the EAGLE atopic dermatitis (eczema) GWAS (Paternoster et al., 2015), and for each signal we defined a strict LD-based interval thresholded at r^2^ > 0.6. Among 94 Mendelian disease genes overlapping the intervals, 15 are associated with the general “Abnormality of the skin” HPO term. On further investigation, one of these had evidence of eQTL colocalisation (Giambartolomei et al., 2014) in the skin - 18q21.33 locus represented by the lead SNP rs35112771 (intronic) with the *p*-value of 2.1 × 10^−5^. The region showed convincing evidence of colocalisation (83.5%) with *SERPINB7* eQTLs only in the sun-unexposed skin tissue type of GTEx study ver. 7 (Ardlie et al., 2015). Crucially, *SERPINB7*’s expression is elevated in the epidermis, and nonsense mutations in the gene cause Nagashima-Type Palmoplantar Keratosis, characterized by well-demarcated diffuse hyperkeratosis with redness on palms and feet (Kubo et al., 2013). This phenotype is widely found in a range of eczematous conditions (Thestrup-Pedersen, Andersen, Menné, & Veien, 2001; J.-H. Lee et al., 2009; Agner et al., 2015) and would suggest that common mutations affecting *SERPINB7* expression could partially explain the missing heritability in atopic dermatitis GWAS.

## Discussion

MendelVar is a unique webserver/pipeline dedicated to automatic assessment of the genetic and phenotypic link between Mendelian disease genes and hits at complex trait GWAS loci. It maps Mendelian disease to relevant variants and disease phenotypes as well as descriptions from a comprehensive sampling of public databases reporting Mendelian gene-disease associations. MendelVar’s rich database contains 4,667 unique genes, 11,412 gene-disease relationships - 83.2% with a disease description, 86% with Human Phenotype Ontology annotation, 62.3% with Disease Ontology annotation; all complemented by ∼100,000 ClinVar rare variants. The Mendelian disease phenotypes and other gene ontology terms are then tested for enrichment in a GWAS-suitable framework. All these features are bundled into one streamlined process beginning with input of genomic coordinates or variant rsIDs by the user. MendelVar delegates to the user a great deal of flexibility regarding workflow – allowing definition of a crude fixed interval around input genomic positions (GRCh38/hg38 or GRCh37/hg19), mapping positions or rsIDs to flexible LD-based regions or input of custom genomic intervals derived using independent methods. Our goal was to integrate all publicly available data sources mapping Mendelian disease to genes, hence inclusion of Orphanet, Genomics England PanelApp and DECIPHER to supplement the chief authority on Mendelian disease - OMIM. This strategy proved beneficial as MendelVar includes 629 novel genes and at least 1,875 new gene-disease associations compared to OMIM. For instance, *CREB3L3* and *CDX1* are not assigned any disorders on OMIM but are on the “Severe hypertriglyceridaemia” panel and “Non-syndromic familial congenital anorectal malformations” panel, respectively, in the Genomics England PanelApp.

Absence of given gene-disease connections in OMIM can be not only because of delay due to its reliance on literature rather than clinical data but also stricter requirements for inclusion and redundancy. Some links presented in the non-OMIM sources and supported by the older literature have been deemed not strong enough to deserve a separate entry in OMIM, e.g. the link between *KCNJ8* and Cantú syndrome (Cooper et al., 2014) reported in DECIPHER or a link between *PLXND1* and Moebius syndrome in Orphanet (Tomas-Roca et al., 2015). Each gene-disease entry in MendelVar results table lists all the data sources supporting the causal link, allowing case-by-case assessment by the user.

Some information is likely to be partially or wholly redundant due to different criteria for assignment to related conditions. For instance, DECIPHER reports *SATB2* to be linked to simply “cleft palate”, whereas in OMIM the gene is mapped to Glass syndrome, one of which hallmarks is cleft palate. Similarly, Orphanet can present alternative assignment of genes to rare diseases relative to OMIM. For instance, *TRDN* is assigned to Romano-Ward syndrome on Orphanet and “Ventricular tachycardia, catecholaminergic polymorphic, 5, with or without muscle weakness” on OMIM, despite the fact that Romano-Ward syndrome is also present in the OMIM database; both diseases are forms of cardiac arrhythmias.

Integration of Mendelian disease genetics into prioritization of genes at GWAS loci may have a positive impact on drug discovery pipelines. Studies have shown that the probability of progression along drug development pipeline is greater for target-indication pairs with genetic evidence, and that OMIM genetic evidence has a greater effect on approval than does GWAS evidence, e.g. phase I to approval progression risk ratio for GWAS=1.4 (95% CI 1.1-1.7), OMIM=2.7 (95% CI 2.4-3.1) (King et al., 2019; Nelson et al., 2015). Thorough analysis has demonstrated that this difference is unlikely to be due to a statistical artefact or reverse causation, whereby the OMIM entry is influenced by the knowledge of successful drug trials (King et al., 2019). However, the higher probability of progression along drug development pipelines for OMIM genes may be influenced by Mendelian conditions having larger genetic risk factor effects than common GWAS traits (rather than just increased likelihood of identifying the true causal gene) and so it remains to be seen whether the use of Mendelian genetic data to link GWAS hits to causal genes will result in the same positive impact on drug approval. Regardless, genes prioritized by MendelVar may make for attractive targets for pharmaceutical companies as they may prove to be effective for more than one condition (i.e. both the common GWAS condition and the rarer Mendelian condition). Examples of successful drugs targeting Mendelian disease genes which are used to treat common conditions include PCSK9 antibodies and orexin antagonists. While monogenic forms of hypercholesterolemia which leads to early development of cardiovascular disease have implicated key genes in the LDL-cholesterol transport pathway, including *PCSK9*, PCSK9 monoclonal antibodies evolocumab and alirocumab are now indicated for treatment of common hyperlipidemia. Dominant loss-of-function mutations in the orexin gene can cause narcolepsy; an orexin receptor antagonist drug (suvorexant) has proven effective for treating insomnia (Coleman, Gotter, Herring, Winrow, & Renger, 2017).

As shown by examples of identification of candidate genes and enriched disease phenotypes/classes in intelligence, male patterned baldness and atopic dermatitis GWAS, we envisage that MendelVar will be especially useful in annotation and interpretation of GWAS results, but flexibility of the user input enables other applications, such as in EWAS (epigenome-wide association studies). Ours and previous analyses (Freund et al., 2018) underscore that MendelVar can be relied on to highlight enrichment of relevant biological processes implicated in investigated trait/disease. However, the degree of enrichment observed can vary depending on the relative contribution of Mendelian disease genes to a given trait.

MendelVar’s structured and comprehensive output allows inclusion into GWAS interpretation pipelines to complement other methods such as: Bayesian variant fine-mapping, variant pathogenicity prediction, molecular QTL mapping and colocalisation, chromosome confirmation capture for identifying regulatory loops. As an example, we present *SERPINB7* as a new putative candidate gene in atopic dermatitis GWAS due to not only a phenotypic match between its target Mendelian disease and eczema but also due to colocalisation evidence in the skin. In another example of usefulness of contrasting MendelVar with other methods, after initial phenotype-matched selection of 6 genes in a cluster of 20 keratin genes at a male-pattern baldness locus, cross-referencing with the FUMA pipeline further halves the number of candidate genes to 3.

In summary, MendelVar provides a useful complementary method aiding candidate gene prioritisation in GWAS. MendelVar highlights disease processes, organ abnormalities and symptoms enriched among Mendelian disease genes which are likely dysregulated in phenotypically-related complex traits.

### Availability

MendelVar webserver is publicly available on https://mendelvar.mrcieu.ac.uk Detailed MendelVar tutorial can be found on https://bitly.com/MendelVar or https://www.notion.so/mendelvar/MendelVar-tutorial-ab91d2a6acb846f2b9f2978fcd942dd5

Code to reproduce MendelVar pipeline: https://github.com/MRCIEU/mendelvar_standalone

## Materials and Methods

### Gene annotations

APPRIS (Rodriguez et al., 2013) isoform definitions were downloaded from its website along with the matching GENCODE GFF annotation files: v19 (for GChr37) and v31 (for GChr38). We used the canonical APPRIS transcripts to define gene coordinates. When more than one canonical isoform was available in APPRIS, we chose the longest transcript. Only 20,738 coding genes are present in APPRIS. Around 37,000 more non-coding genes are annotated in GENCODE - RNA genes, hypothetical genes etc. which do not have a defined CDS. We included those genes in our dataset by selecting the longest transcript available in GENCODE.

### Database input filtering

OMIM filtering was as follows: disease gene mapping key must be 3 (also prefix #), ie. with known molecular basis - known causal gene. Gene-disease relationships reported in DECIPHER DDG2P were kept if they were in category: “probable” or “confirmed” but not “possible”.

Data from DECIPHER and Orphanet required additional cleaning, in terms of checking presence of the disease/gene MIMs in the OMIM database and their up-to-date status – some MIM IDs needed upgrading from deprecated to current IDs. We also included checks to establish congruence between MIM gene IDs and gene symbol and complete missing MIM gene IDs.

### Integration of OMIM, Orphanet, DECIPHER and Genomics England PanelApp

The four different data sources for Mendelian disease-gene mapping (OMIM, Orphanet, DECIPHER, Genomics Panel App) were subsequently integrated. Genes were all standardised by HGNC ID (Eyre et al., 2006), Ensembl IDs, and HGNC approved symbol (in that order), and disease names through OMIM IDs, Orphanet IDs and OMIM names.

### ClinVar variant data processing

Variants downloaded from ClinVar FTP in the variant_summary.txt.gz file were filtered to retain only those that contain “pathogenic", “likely pathogenic” or “risk factor” among their effects. Variants spanning many genes were eliminated, to keep variants directly linked to a single or a very small number of genes. Variants missing coordinates were also discarded, and phenotype descriptions matched to MIM IDs whenever possible. This filtering strategy resulted in retention of approximately 20% of variants – 105,824 and 104,099 in GRCh37 and GRCh38, respectively.

### Ontology processing

For multi-level ontologies – DO, HPO, GO, we propagated all the transitive “is_a”, “part_of” relationships up to the root using the R package ontologyIndex (Greene, Richardson, & Turro, 2016), and included all the parent terms in addition to leaf terms, exclusive of the root terms in each ontology, because these are uninformative.

We eliminated all Inferred from Electronic Annotations (IEA) and entries with “NOT” qualifier (ie. gene is NOT characterised by the term), as IEA annotations are not manually curated, often inferred just by text mining algorithms and can be thus unreliable.

We only retained the “Phenotypic abnormality” HPO ontology and discarded children of HP:0000005 Mode of inheritance, HP:0031797 Clinical course, HP:0040279 Frequency, HP:0012823 Clinical modifier as these encompass a small number of child terms and do not correlate with the mechanistic basis of disease. HPO slim ontology was created by pruning the tree to only 25 direct descendants of the root HP:0000118 Phenotypic abnormality term.

In REACTOME, we used the Ensembl2Reactome_All_Levels.txt file which contains gene annotations across all pathway levels.

Finally, we subsetted all the ontologies only to the Mendelian disease genes in the MendelVar database and eliminated ontology annotations of the genes with no evidence for disease causality, as we want to test for term enrichment relative to genes linked to Mendelian disease rather any gene as background.

### Enrichment testing with INRICH

INRICH was revealed to be one of the most sensitive and specific methods for conducting overrepresentation analysis on GWAS data (De Leeuw, Neale, Heskes, & Posthuma, 2016) and is a fast C++ software, which makes it ideal for our application.

INRICH takes a list of associated genomic intervals and tests for enrichment against gene sets. The intervals need not be independent from each other, as overlapping intervals are merged, as well as overlapping genes belonging to the same gene set (P. H. Lee et al., 2012). Background interval sets for enrichment testing are permuted to match the input set in terms of the number of SNPs, SNP density and number of overlapping genes which removes a lot of bias when using GWAS-derived genomic intervals.

## Acknowledgements

MKS, TRG and LP work in a research unit funded by the UK Medical Research Council (MC_UU_00011/4). MKS is funded by an Academy of Medical Sciences Springboard Award (awarded to LP), which is supported by the Wellcome Trust, The Government Department for Business, Energy and Industrial Strategy, the Global Challenges Research Fund and the British Heart Foundation [SBF003\1094].

## Resources

Gencode:

*GRCh38/hg38*

ftp://ftp.ebi.ac.uk/pub/databases/gencode/Gencode_human/release_31/gencode.v31.annotation.gff3.gz

*GRCh37/hg19*

ftp://ftp.ebi.ac.uk/pub/databases/gencode/Gencode_human/release_19/gencode.v19.annotation.gff3.gz

HUGO Gene Nomenclature Committee (HGNC)

ftp://ftp.ebi.ac.uk/pub/databases/genenames/new/tsv/hgnc_complete_set.txt

APPRIS isoforms

*For version hg19 (matched to Gencode version 19)*

http://apprisws.bioinfo.cnio.es/pub/current_release/datafiles/homo_sapiens/GRCh37/appris_data.principal.txt

*For version hg38 (matched to Gencode version 31)*

http://apprisws.bioinfo.cnio.es/pub/current_release/datafiles/homo_sapiens/GRCh38/appris_data.principal.txt

1000 Genomes

*GRCh37/hg19*

http://ftp.1000genomes.ebi.ac.uk/vol1/ftp/release/20130502/

*GRCh38/hg38*

http://ftp.1000genomes.ebi.ac.uk/vol1/ftp/release/20130502/supporting/GRCh38_positions

HapMap II recombination hotspots

ftp://ftp.ncbi.nlm.nih.gov/hapmap/recombination/2011-01_phaseII_B37/genetic_map_HapMapII_GRCh37.tar.gz

LDlink LDproxy

https://ldlink.nci.nih.gov/?tab=ldproxy

UCSC liftOver

https://genome.ucsc.edu/cgi-bin/hgLiftOver

Giggle

https://github.com/ryanlayer/giggle

INRICH

https://atgu.mgh.harvard.edu/inrich/

OMIM

https://www.omim.org/api

Orphanet

https://www.orpha.net/

*Rare diseases and cross referencing*

http://www.orphadata.org/data/xml/en_product1.xml

*Linearisation of disorders*

http://www.orphadata.org/data/xml/en_product7.xml

*Rare diseases with their associated genes*

http://www.orphadata.org/data/xml/en_product6.xml

Genomics England

https://panelapp.genomicsengland.co.uk/api/v1/

DECIPHER

https://decipher.sanger.ac.uk/

http://www.ebi.ac.uk/gene2phenotype/downloads/DDG2P.csv.gz

ClinVar

https://www.ncbi.nlm.nih.gov/clinvar/

ftp://ftp.ncbi.nlm.nih.gov/pub/clinvar/tab_delimited/variant_summary.txt.gz

Disease Ontology

http://disease-ontology.org/

*Full ontology*

https://raw.githubusercontent.com/DiseaseOntology/HumanDiseaseOntology/master/src/ontology/doid-merged.obo

*Slim ontology*

https://raw.githubusercontent.com/DiseaseOntology/HumanDiseaseOntology/master/src/ontology/subsets/DO_AGR_slim.obo

Human Phenotype Ontology

https://hpo.jax.org/

*Ontology*

https://raw.githubusercontent.com/obophenotype/human-phenotype-ontology/master/hp.obo

*Annotation*

http://compbio.charite.de/jenkins/job/hpo.annotations.current/lastSuccessfulBuild/artifact/current/phenotype.hpoa

Freund et al. gene sets

https://github.com/bogdanlab/gene_sets/tree/master/mendelian_gene_sets

Gene Ontology

*Full ontology*

http://current.geneontology.org/ontology/go-basic.obo

*Slim ontology*

http://current.geneontology.org/ontology/subsets/goslim_generic.obo

*Annotation*

ftp://ftp.ebi.ac.uk/pub/databases/GO/goa/HUMAN/goa_human.gaf.gz

ConsensusPathDB

http://consensuspathdb.org/

Pathway Commons

https://www.pathwaycommons.org/

https://www.pathwaycommons.org/archives/PC2/v12/PathwayCommons12.All.hgnc.gmt.gz

Reactome

https://reactome.org/

https://reactome.org/download/current/Ensembl2Reactome_All_Levels.txt

