## Supplementary Figures and Tables for "MendelVar: gene prioritization at GWAS loci using phenotypic enrichment of Mendelian disease genes"

#### Supplementary Figures and Tables captions

**Figure S1. Top enrichment results for Disease Ontology (DO) terms among Mendelian disease genes located within 1 Mbp of lead SNPs in Savage et al. (2018) intelligence GWAS.**

**Figure S2. Top enrichment results for Pathway Commons terms among Mendelian disease genes located within 1 Mbp of lead SNPs in Yap et al. (2018) male pattern baldness GWAS.**

**Figure S3. Top enrichment results for ConsensusPathDb terms among Mendelian disease genes located within 1 Mbp of lead SNPs in Yap et al. (2018) male pattern baldness GWAS.**

**Figure S4. Top enrichment results for Human Phenotype Ontology (HPO) terms among Mendelian disease genes located within 1 Mbp of lead SNPs in Yap et al. (2018) male pattern baldness GWAS.**

**Table S1. Sample truncated output from the MendelVar disease gene overlap table.** Each row (here transposed) in the table represents a single Mendelian disease gene overlapping our test interval.

**Table S2. Sample truncated output from the MendelVar variant overlap table.** Each row (here transposed) in the table represents single ClinVar pathogenic/likely pathogenic/risk variant overlapping our test interval.

**Table S3. Comparison of MendelVar with related Mendelian disease- and enrichment- centred annotation tools.**

### Supplementary Figures and Tables

#### Figure S1

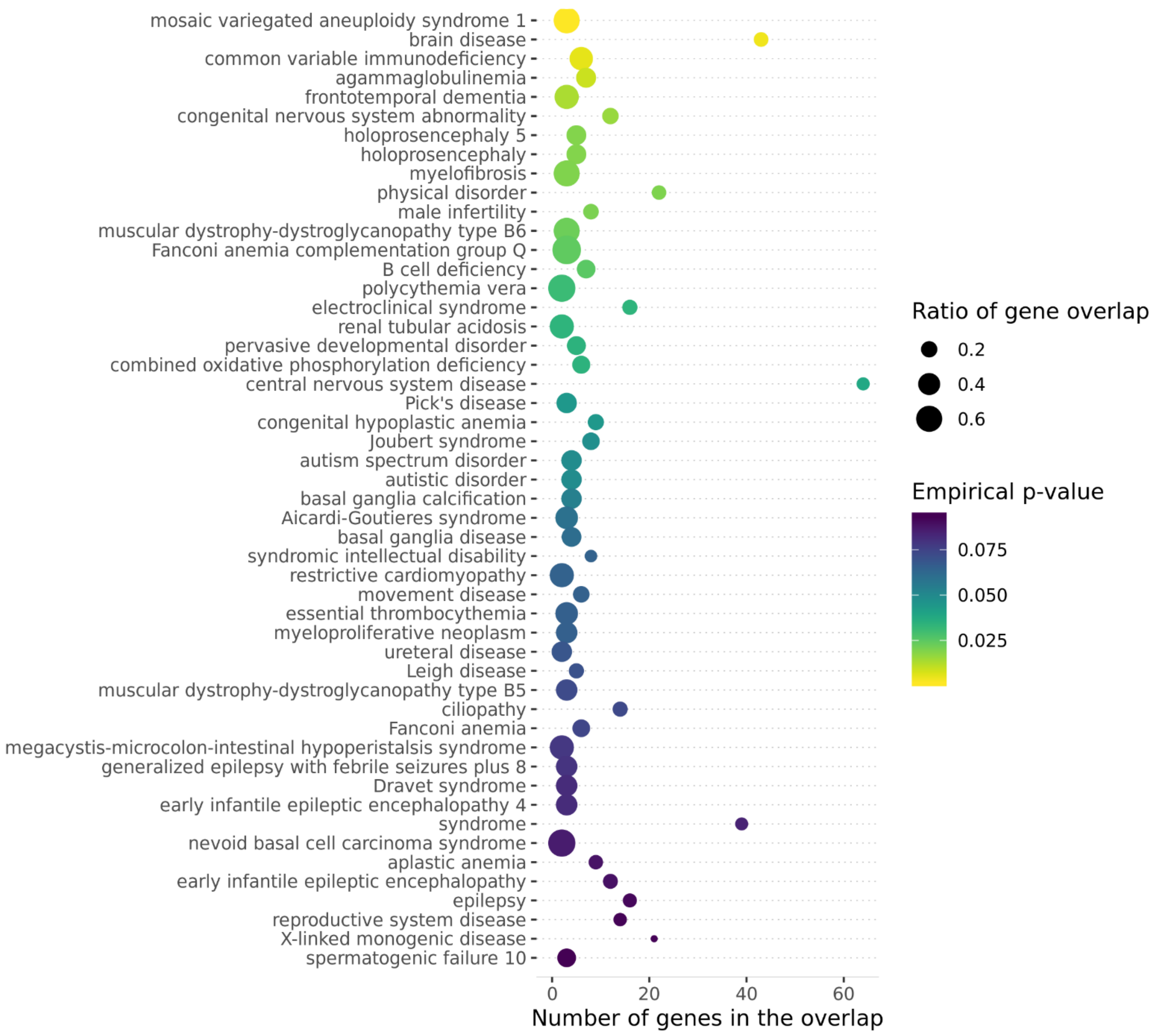

**Figure S2**

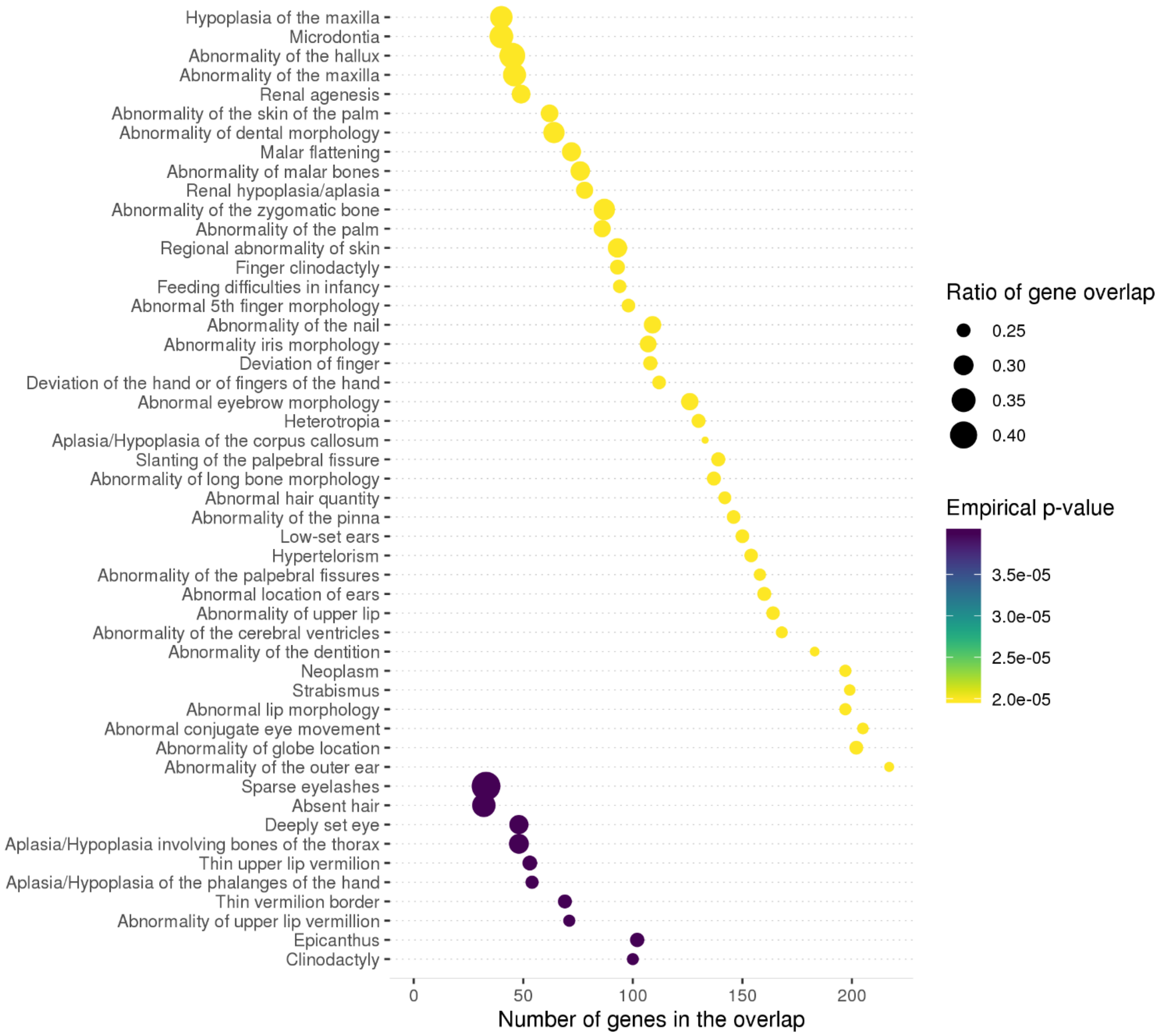
Hypoplasia of the maxilla
Microdontia
Abnormality of the hallux
Abnormality of the maxilla
Renal agenesis
Abnormality of the skin of the palm
Abnormality of dental morphology
Malar flattening
Abnormality of malar bones
Renal hypoplasia/aplasia
Abnormality of the zygomatic bone
Abnormality of the palm
Regional abnormality of skin
Finger clinodactyly
Feeding difficulties in infancy
Abnormal 5th finger morphology
Abnormality of the nail
Abnormality iris morphology
Deviation of finger
Deviation of the hand or of fingers of the hand
Abnormal eyebrow morphology
Heterotropia
Aplasia/Hypoplasia of the corpus callosum
Slanting of the palpebral fissure
Abnormality of long bone morphology
Abnormal hair quantity
Abnormality of the pinna
Low-set ears
Hypertelorism
Abnormality of the palpebral fissures
Abnormal location of ears
Abnormality of upper lip
Abnormality of the cerebral ventricles
Abnormality of the dentition
Neoplasm
Strabismus
Abnormal lip morphology
Abnormal conjugate eye movement
Abnormality of globe location
Abnormality of the outer ear
Sparse eyelashes
Absent hair
Deeply set eye
Aplasia/Hypoplasia involving bones of the thorax
Thin upper lip vermilion
Aplasia/Hypoplasia of the phalanges of the hand
Thin vermilion border
Abnormality of upper lip vermillion
Epicanthus
Clinodactyly
0
50
100
150
200
Number of genes in the overlap
Ratio of gene overlap
0.25
0.30
0.35
0.40
Empirical p-value
3.5e-05
3.0e-05
2.5e-05
2.0e-05

**Figure S3**

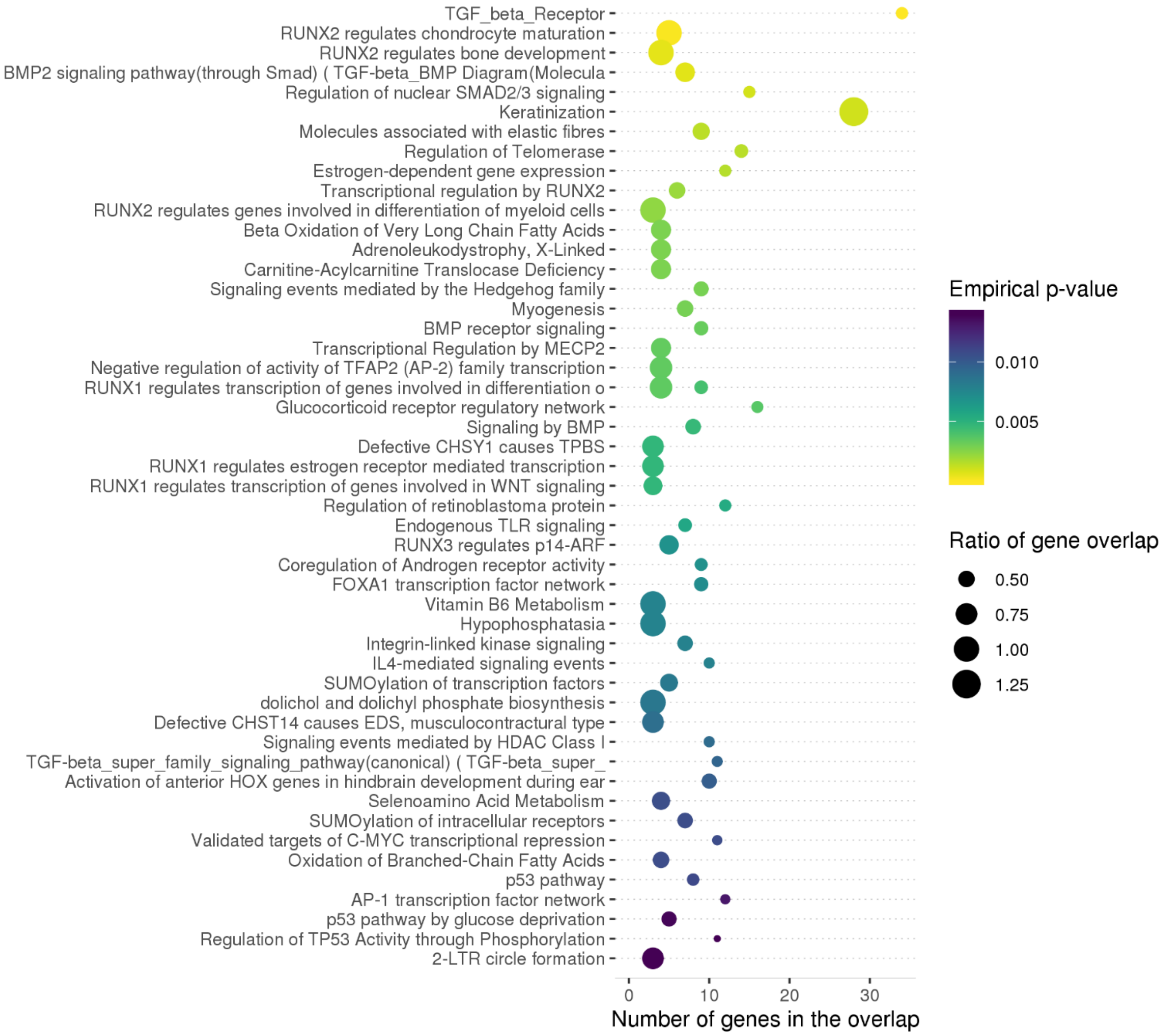
TGF_beta_Receptor
RUNX2 regulates chondrocyte maturation
RUNX2 regulates bone development
BMP2 signaling pathway(through Smad) ( TGF-beta_BMP Diagram(Molecula
Regulation of nuclear SMAD2/3 signaling
Keratinization
Molecules associated with elastic fibres
Regulation of Telomerase
Estrogen-dependent gene expression
Transcriptional regulation by RUNX2
RUNX2 regulates genes involved in differentiation of myeloid cells
Beta Oxidation of Very Long Chain Fatty Acids
Adrenoleukodystrophy, X-Linked
Carnitine-Acylcarnitine Translocase Deficiency
Signaling events mediated by the Hedgehog family
Myogenesis
BMP receptor signaling
Transcriptional Regulation by MECP2
Negative regulation of activity of TFAP2 (AP-2) family transcription
RUNX1 regulates transcription of genes involved in differentiation o
Glucocorticoid receptor regulatory network
Signaling by BMP
Defective CHSY1 causes TPBS
RUNX1 regulates estrogen receptor mediated transcription
RUNX1 regulates transcription of genes involved in WNT signaling
Regulation of retinoblastoma protein
Endogenous TLR signaling
RUNX3 regulates p14-ARF
Coregulation of Androgen receptor activity
FOXA1 transcription factor network
Vitamin B6 Metabolism
Hypophosphatasia
Integrin-linked kinase signaling
IL4-mediated signaling events
SUMOylation of transcription factors
dolichol and dolichyl phosphate biosynthesis
Defective CHST14 causes EDS, musculocontractural type
Signaling events mediated by HDAC Class I
TGF-beta_super_family_signaling_pathway(canonical) ( TGF-beta_super_
Activation of anterior HOX genes in hindbrain development during ear
Selenoamino Acid Metabolism
SUMOylation of intracellular receptors
Validated targets of C-MYC transcriptional repression
Oxidation of Branched-Chain Fatty Acids
p53 pathway
AP-1 transcription factor network
p53 pathway by glucose deprivation
Regulation of TP53 Activity through Phosphorylation
2-LTR circle formation
0
10
20
30
Number of genes in the overlap
Empirical p-value
0.010
0.005
Ratio of gene overlap
0.50
0.75
1.00
1.25

**Figure S4**

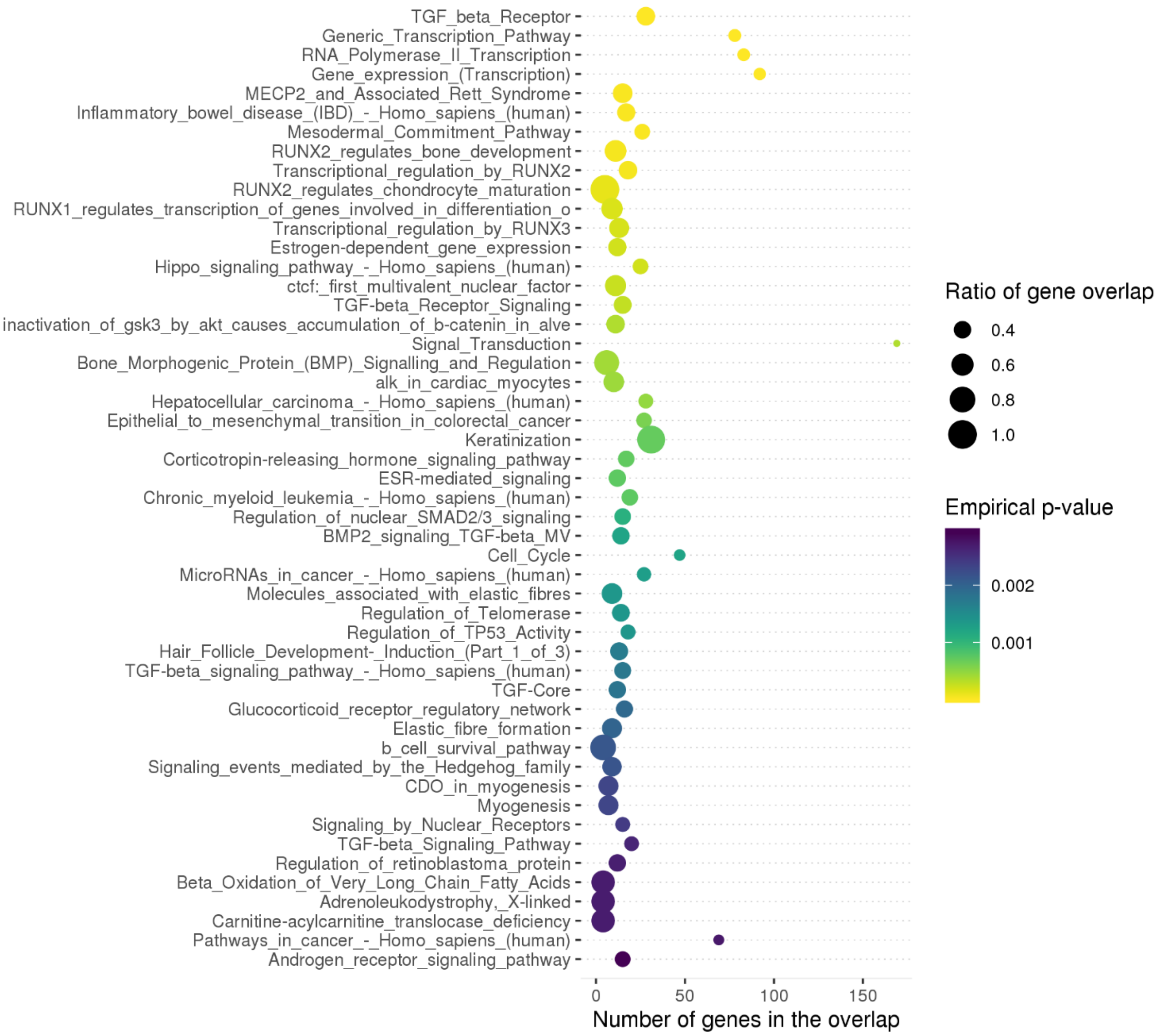

TGF_beta_Receptor
Generic_Transcription_Pathway
RNA_Polymerase_II_Transcription
Gene_expression_(Transcription)
MECP2_and_Associated_Rett_Syndrome
Inflammatory_bowel_disease_(IBD)_-_Homo_sapiens_(human)
Mesodermal_Commitment_Pathway
RUNX2_regulates_bone_development
Transcriptional_regulation_by_RUNX2
RUNX2_regulates_chondrocyte_maturation
RUNX1_regulates_transcription_of_genes_involved_in_differentiation_o
Transcriptional_regulation_by_RUNX3
Estrogen-dependent_gene_expression
Hippo_signaling_pathway_-_Homo_sapiens_(human)
ctcf:_first_multivalent_nuclear_factor
TGF-beta_Receptor_Signaling
inactivation_of_gsk3_by_akt_causes_accumulation_of_b-catenin_in_alve
Signal_Transduction
Bone_Morphogenic_Protein_(BMP)_Signalling_and_Regulation
alk_in_cardiac_myocytes
Hepatocellular_carcinoma_-_Homo_sapiens_(human)
Epithelial_to_mesenchymal_transition_in_colorectal_cancer
Keratinization
Corticotropin-releasing_hormone_signaling_pathway
ESR-mediated_signaling
Chronic_myeloid_leukemia_-_Homo_sapiens_(human)
Regulation_of_nuclear_SMAD2/3_signaling
BMP2_signaling_TGF-beta_MV
Cell_Cycle
MicroRNAs_in_cancer_-_Homo_sapiens_(human)
Molecules_associated_with_elastic_fibres
Regulation_of_Telomerase
Regulation_of_TP53_Activity
Hair_Follicle_Development-_Induction_(Part_1_of_3)
TGF-beta_signaling_pathway_-_Homo_sapiens_(human)
TGF-Core
Glucocorticoid_receptor_regulatory_network
Elastic_fibre_formation
b_cell_survival_pathway
Signaling_events_mediated_by_the_Hedgehog_family
CDO_in_myogenesis
Myogenesis
Signaling_by_Nuclear_Receptors
TGF-beta_Signaling_Pathway
Regulation_of_retinoblastoma_protein
Beta_Oxidation_of_Very_Long_Chain_Fatty_Acids
Adrenoleukodystrophy,_X-linked
Carnitine-acylcarnitine_translocase_deficiency
Pathways_in_cancer_-_Homo_sapiens_(human)
Androgen_receptor_signaling_pathway
0
50
100
150
Number of genes in the overlap
Ratio of gene overlap
0.4
0.6
0.8
1.0
Empirical p-value
0.002
0.001

**Table S1**

| | Example |
|---|---|
| **ID** | rs80293268 |
| **chrom** | chr1 |
| **interval_start** | 7647519 |
| **interval_end** | 8647519 |
| **gene_name** | CAMTA1 |
| **alias_symbol** | KIAA0833 |
| **hgnc_id** | 18806 |
| **Ensembl_gene_id** | ENSG00000171735 |
| **gene_start** | 6785454 |
| **gene_end** | 7769706 |
| **strand** | + |
| **omim_gene_id** | 611501 |
| **omim_disease_id** | 614756 |
| **orphanet_id** | 314647 |
| **disease_name** | Cerebellar ataxia, nonprogressive, with mental retardation |
| **hpo** | 0000752;0000718;0002120;0006919;0001263;0000276;0001260;0002317;0002019;0100540;0000307;0000490;0001348;0002403;0000414;0001319;0000160;0001321;0000639;0010867;0025517;0001249;0002003;0000445;0002470;0002536;0000369;0001256;0001310;0001250;0000729;0000494;0001251;0002080;0400005;0000179;0012433;0002275;0000343;0011067;0025191;0002354;0000750;0000337;0410170;0007256;0000463;0000316;0002020;0000256;0000411;0000486 |
| **do** | 50998 |
| **source** | Decipher;Genomics England;OMIM;Orphanet |
| **omim_link** | https://www.omim.org/entry/614756 |

| | |
|---|---|
| **disease_description** | Nonprogressive cerebellar ataxia with mental retardation is an autosomal dominant neurodevelopmental disorder characterized by mildly delayed psychomotor development, early onset of cerebellar ataxia, and intellectual disability later in childhood and adult life. Other features may include neonatal hypotonia, dysarthria, and dysmetria. Brain imaging in some patients shows cerebellar atrophy. Dysmorphic facial features are variable (summary by Thevenon et al., 2012). |

**Table S2**

| | Example |
|---|---|
| **ID** | rs80293268 |
| **chrom** | chr1 |
| **interval_start** | 7647519 |
| **interval_end** | 8647519 |
| **dbsnp_dbvar_id** | rs1135401818 |
| **variant_start** | 7677682 |
| **variant_end** | 7677682 |
| **cyto_location** | 1p36.23 |
| **ref_allele** | C |
| **alt_allele** | T |
| **effect** | Likely pathogenic |
| **HGSV_notation** | NM_015215.4(CAMTA1):c.2863C>T (p.Arg955Trp) |
| **gene** | CAMTA1 |
| **disease_name(s)** | Cerebellar ataxia, nonprogressive, with mental retardation |
| **disease_omim** | 614756 |
| **disease_orphanet** | NA |
| **clinvar_link** | http://www.ncbi.nlm.nih.gov/clinvar/variation/431151 |
| **VCV** | 431151 |
| **RCV** | 496160 |
| **allele_id** | 424639 |
| **quality_rating** | criteria provided, single submitter |

**Table S3**

| | MendelVar | MARRVEL | VarfromPDB | DisGeNET | FUMA | clusterProfiler | DAVID | Enrichr |
|---|---|---|---|---|---|---|---|---|
| *Purpose* | Annotation and gene prioritization at GWAS loci using phenotypic enrichment of Mendelian disease genes | Cross-displinary integration of genetic and molecular data from human and model organisms for exploration of genes and variants | Mining genes and variants related to a Mendelian disorder from multiple public curated databases and literature | Integrating data from curated repositories, GWAS catalogues, animal models and scientific literature to support versatile research | Annotation of variants and genes at GWAS loci using multiple resources with interpretation aided by interactive visualisation and gene/tissue prioritisation | Flexible pathway/gene set enrichment and visualisation across many species for high-throughput genomics research | Gene set annotation, enrichment and clustering analysis with curated libraries for high-throughput genomics research | Comprehensive gene set enrichment analysis with curated libraries aided by interactive visualisation for high-throughput genomics research |
| *Release type* | Webserver | Webserver | R package requiring custom downloads | Webserver, R package, SQLite database | Webserver | R package | Webserver | Webserver, R package |
| *Type of input accepted* | List of genomic coordinates: ranges, single positions, variant rsIDs | A single gene name, mutation (coordinate or HGSV) or both | Disease name, phenotype keywords | List of gene, disease names/IDs or variant rsIDs | GWAS full summary stats (+ genomic range coordinates), or list of gene IDs | List of gene IDs/names | List of gene IDs/names | List of gene IDs/names or genomic coordinates |
| *Mapping of genes to Mendelian disease* | Yes | Yes | Yes | Yes | Yes | No | Yes | Yes |
| *Mendelian disease databases included* | OMIM, Orphanet, Genomics England, DECIPHER | OMIM | OMIM, Orphanet | OMIM, Orphanet, Genomics England | OMIM | No | OMIM | OMIM, DisGeNET, ClinVar |
| *Mendelian disease descriptions included* | Yes | No | No | No | No | No | No | No |
| *Querying for clinically established, Mendelian disease mutations* | Prefiltered for disease mutations - ClinVar | Unfiltered - ClinVar, Geno2MP | Unfiltered - ClinVar, Humsavar | ClinVar, HumsaVar | No | No | No | No |
| *Ontology enrichment testing* | Yes | No | No | No | Yes | Yes | Yes | Yes |
| *Enrichment testing suitable for genomic input (e.g. GWAS)?** | Yes | No | No | No | No | No | No | No |
| *Human disease ontologies included* | Human Phenotype Ontology, Disease Ontology, Freund et al. (2018) | Human Phenotype Ontology (via Geno2MP) | Human Phenotype Ontology | Human Phenotype Ontology and Disease Ontology (high level terms) | No | Disease Ontology | No | Human Phenotype Ontology |
| *Other ontologies / pathways included* | Gene Ontology, Reactome, Pathway Commons, ConsensusPathDb | Gene Ontology | No | No | Gene Ontology, MsigDB, WikiPathways | Gene Ontology, KEGG Pathway, Reactome, MsigDB, WikiPathways | Gene Ontology, KEGG Pathway, Reactome, Biocarta and others | Gene Ontology, KEGG Pathway, Reactome, Biocarta, WikiPathways and others |

*adjusting for confounders due to genomic location with parametrised re-sampling to generate background set
